# Programming mRNA decay to modulate synthetic circuit resource allocation

**DOI:** 10.1101/079939

**Authors:** Ophelia S. Venturelli, Mika Tei, Stefan Bauer, Leanne Jade G. Chan, Christopher J. Petzold, Adam P. Arkin

## Abstract

Synthetic biomolecular networks embedded in host-cells compete with cellular processes for limited intracellular resources. Resource utilization is a major variable that impacts synthetic circuit behavior. Here we show that intracellular resources could be diverted from cellular operations to a synthetic circuit by programming global mRNA decay using the sequence-dependent endoribonuclease MazF. Synthetic circuit genes were protected from MazF activity by recoding the gene sequence to eliminate recognition sites, while preserving the amino acid sequence. The expression of a protected fluorescent reporter and the metabolic flux of a high-value metabolite gluconate were significantly enhanced using this genome-scale control strategy. Proteomics measurements discovered a key host translation factor in need of protection to optimize the resource redistribution activity. A dynamic computational model demonstrated that the MazF mRNA-decay feedback loop achieved proportional control of MazF levels in an optimal operating regime. RNA-seq time-series measurements of MazF-induced cells elucidated the dynamic shifts in the transcript abundance and discovered regulatory design elements that could be used to expand the control mechanisms of the MazF resource allocator. Together, these results demonstrated that manipulation of resource allocation is a tunable parameter that can be used to redirect resources away from cellular processes to synthetic circuits to enhance target functions.

## INTRODUCTION

Engineered biological systems have diverse applications in medicine, bioenergy, and agriculture^1^. Novel cellular behaviors can be programmed by interacting networks of biomolecules to process information from the environment and execute target functions. These synthetic biomolecular circuits interact with endogenous cellular processes through competition over shared resources that include ribosomes, tRNAs, polymerases, amino acids, and nucleotides^2^. Resource utilization influences the predictability, function, and evolutionary stability of engineered networks and constrains the achievable parameter space for synthetic circuit design^3^.

Cells operate with a limited resource quota, which produces a trade-off in the partitioning of energy between cellular processes and synthetic circuit functions^1^,^2^,^4^,^5^. A core challenge is to rewire cellular regulation to optimally distribute resources between the host-cell and synthetic circuit processes. While there are numerous mechanisms to control target gene expression including engineered promoters^6^, protein degradation^7^ or CRISPRi^8–10^, limited technologies exist to globally redistribute resources and re-program cellular state. Novel strategies should be developed to manipulate genome-wide gene expression patterns to optimize a target function.

RNA degradation more rapidly and efficiently redistributes ribosomes, a crucial limiting resource in *E. coli*^5^,^11^, compared to transcriptional control. Viruses capitalize on mRNA decay to reduce competition for the host-cell translational machinery during developmental transitions and implement temporal gene expression programs^12^,^13^. To exploit RNA decay for synthetic circuit resource redistribution in *E. coli*, we repurposed a sequence-specific ribonuclease MazF whose recognition site ‘ACA’ is present in 96% of *E. coli* coding sequences. The MazF recognition site can be eliminated from the synthetic circuit while preserving the amino acid content, allowing cellular resources to be re-allocated towards synthetic gene expression by eliminating nearly all competing processes.

Here we show that the MazF resource allocator controllably redistributed core cellular subsystems to support a synthetic circuit and an engineered metabolic pathway. The former was further enhanced by protection of specific host-cell factors and use of the orthogonal RNA polymerase from T7 to transcribe genes in the synthetic circuit. Shotgun proteomics measurements were used to identify a host factor in need of protection to prevent loss of translational efficiency following MazF induction. Our results demonstrated that an mRNA-decay feedback loop was a critical design element for the resource allocator and a dynamic computational model was used to explore the role of feedback in this system. RNA-seq measurements elucidated the relationship between the number of MazF sites and the decay rate of the transcriptome and provided insights into the physiological impact of MazF expression. Further, these data pinpointed regulatory sequences that respond to changes in MazF activity and could be used to expand the control mechanisms of the MazF resource allocator. In sum, these results suggest a platform for global manipulation of resource pools as a key parameter for modulating synthetic circuit behavior.

## RESULTS

### Characterization of inducible MazF for resource allocator design

To explore whether manipulation of resource allocation could predictably modulate circuit behavior, we needed to develop a comprehensive reallocation mechanism that preserved core processes required for a target function while downregulating competing pathways. MazF is a sequence-dependent and ribosome-independent endoribonuclease that cleaves the recognition site ‘ACA’ in single-stranded RNA^14^. 96% of *E. coli* coding sequences contain at least one MazF recognition site (Supplementary Fig. 1). Thus, induction of MazF should inhibit cellular processes other than those protected from its action.

We characterized the impact of MazF on expression of a target gene *mCherry* that contained 9 recognition sites in the coding sequence (*mCherry-U*) or was recoded to not contain any sites using alternative codons (*mCherry-P*). *mazF* was introduced into an intergenic genomic site under control of an aTc-inducible promoter (P_TET_) in an *E. coli* strain deleted for *mazF* (see Materials and Methods). mCherry-P and mCherry-U were expressed at similar levels in the absence of MazF, indicating that recoding the transcript did not significantly modify expression levels (Figure 1b). The MazF induction ratio, defined as the fold change of mCherry-P expression in the presence to absence of MazF, is a metric used to quantify resource redistribution activity. Following 10 hr of induction with 0 or 5 ng/ml aTc, the MazF induction ratio was <1 for mCherry-U and 5 for mCherry-P (Figure 1c). The sequence protection ratio, defined as the ratio of mCherry-P to mCherry-U, was approximately 1 or 19 in the absence or presence of MazF (Figure 1d). Together, these data show that MazF significantly enhanced protected and inhibited unprotected gene expression.

**Figure 1.**
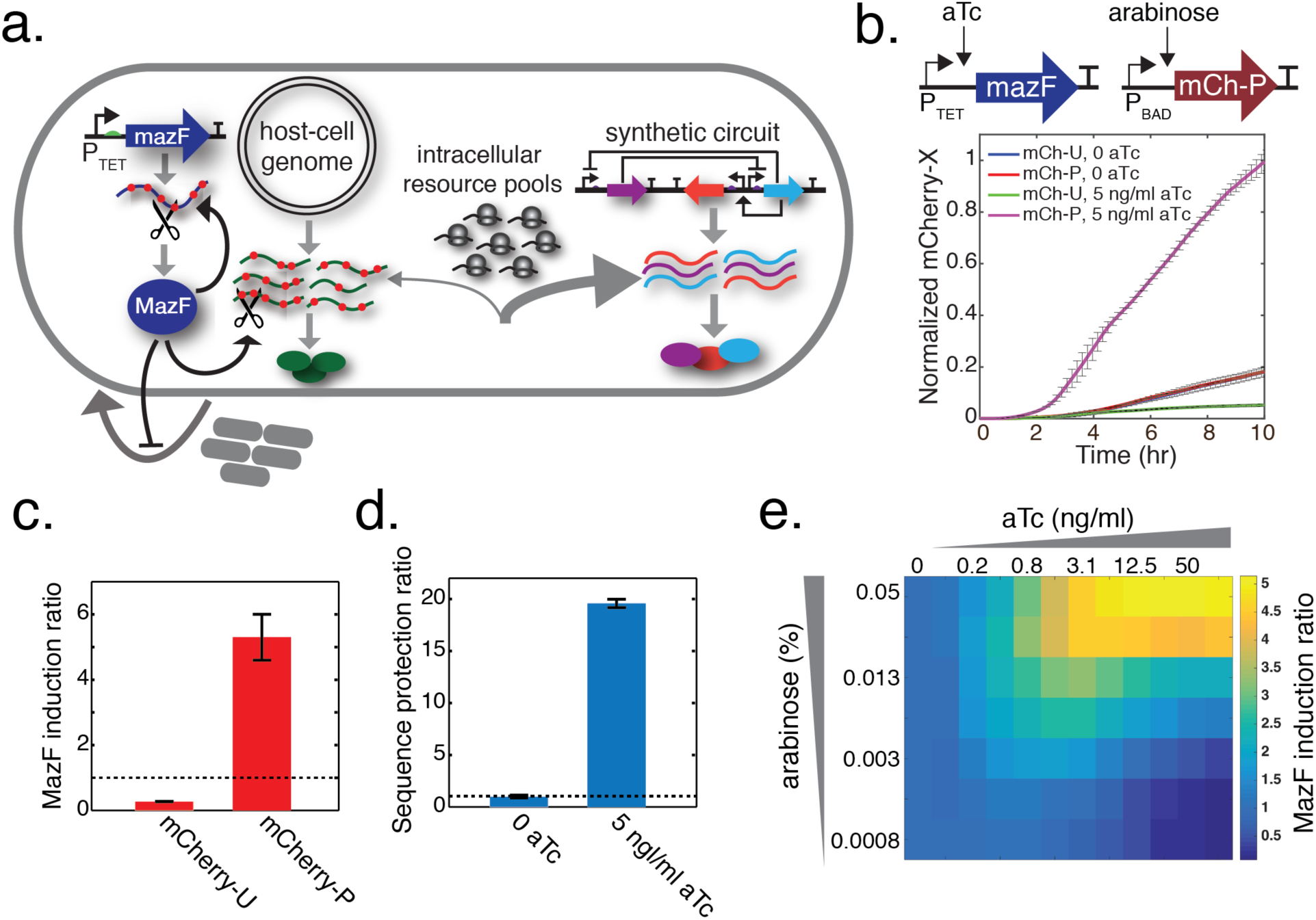
Programming mRNA decay to redistribute resources in *E. coli* to modulate the behavior of synthetic circuits. **(a)** Schematic diagram of the MazF resource allocator. Host-cell transcripts containing MazF recognition sites (‘ACA’) are targeted for cleavage. The MazF site can be removed from target genes while preserving the amino acid sequence. As such, MazF down-regulates transcripts that compete with the protected synthetic circuit for limiting resources, yielding an increase in protected gene protein synthesis. **(b)** MazF and protected mCherry (*mCherry-P*) were controlled by an arabinose or aTc-inducible promoter (top). Time-series measurements of total fluorescence normalized to the maximum steady-state value across conditions as a function of time for cell populations expressing unprotected mCherry (*mCherry-U*) or mCherry-P in the presence (5 ng/ml) of absence (0 ng/ml) of MazF. Cells were induced with 0.05% arabinose. Error bars represent 1 s.d. (n = 3). **(c)** MazF induction ratio, defined as the ratio of expression of mCherry-X in the presence (5 ng/ml aTc) to absence (0 ng/ml aTc) of MazF following induction for 10 hr. Error bars represent 1 s.d. (n = 3). **(d)** Sequence protection ratio, defined as the ratio of expression of mCherry-P to mCherry-U in the presence (5 ng/ml aTc) or absence (0 ng/ml aTc) of MazF. Cells were induced for 10 hr. Error bars represent 1 s.d. (n = 3). **(e)** Heat-map of total fluorescence following 10 hr of induction across a range of arabinose and aTc concentrations.

To map the relationship between MazF expression and resource redistribution activity, growth and mCherry-X (X denotes U or P) expression were measured across a broad range of aTc concentrations. The expression of mCherry-U driven by an arabinose-inducible promoter (P_BAD_) was reduced up to 4-fold in response to aTc (Supplementary Figure 2). The MazF induction ratio of total fluorescence increased (Figure 1e), whereas the total biomass (OD600) decreased as a function aTc (Supplementary Figure 3a). The MazF induction ratio of mCherry fluorescence normalized by OD600 similarly increased with aTc and arabinose, indicating that the biomass normalization factor did not alter the qualitative relationship between MazF activity and mCherry-P expression (Supplementary Figure 3b). These data highlight that mCherry-P expression and biomass synthesis were inversely correlated in response to MazF.

**Figure 2.**
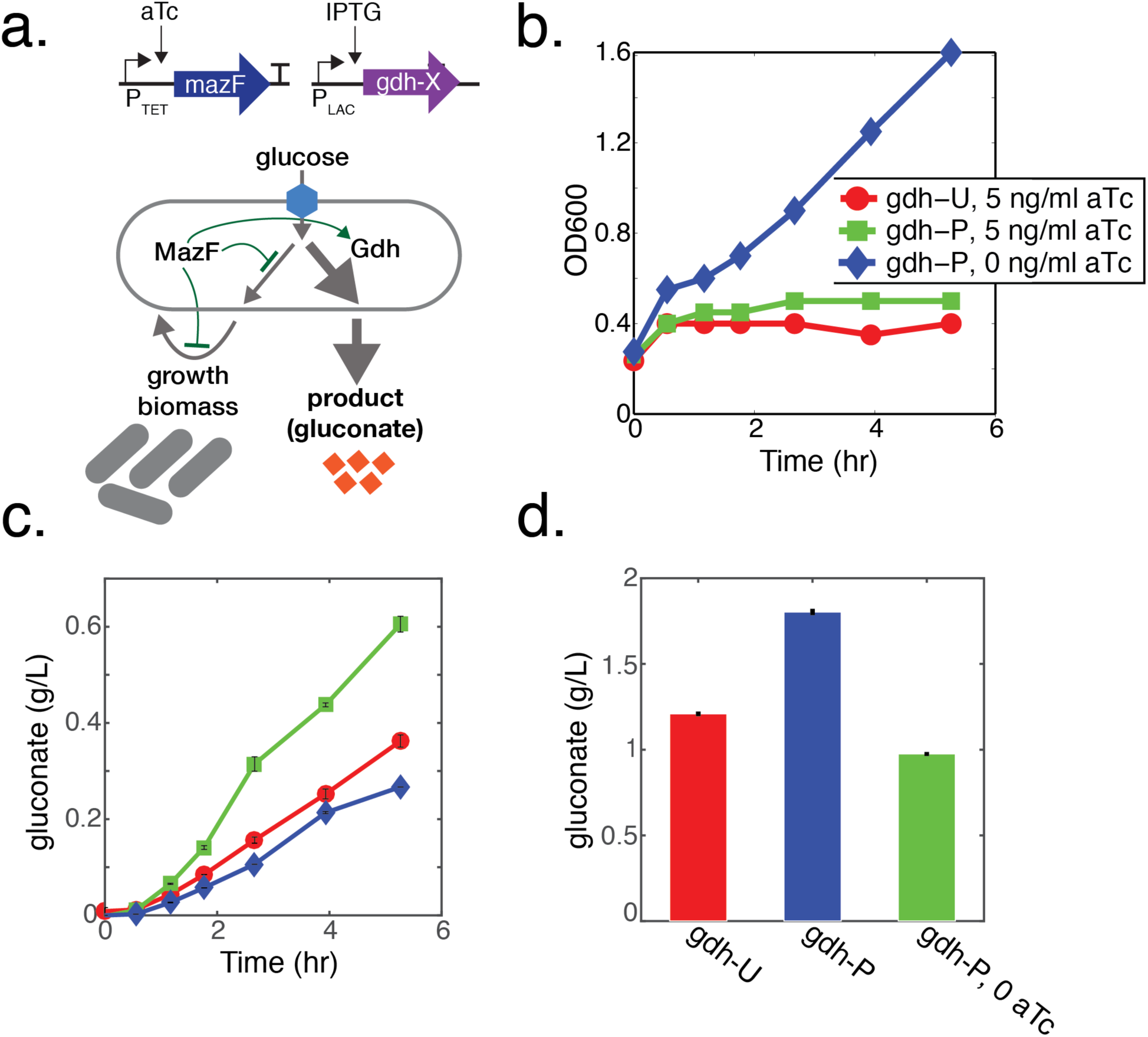
The MazF resource allocator enhanced gluconate production. **(a)** Schematic diagram of the circuit design (top) and gluconate metabolic pathway (bottom). Gdh transforms glucose into gluconate and competes directly with biomass synthesis. MazF and glucose dehydrogenase (gdh) were controlled by an aTc (P_TET_) or IPTG inducible (P_LAC_) promoter. **(b)** Schematic of gluconate circuit (top). OD600 as a function of time for cells expressing Gdh that contained 11 (Gdh-U) or 0 recognition sites (Gdh-P) in response to 5 ng/ml or 0 ng/ml aTc (below). All cultures were induced with 1 mM IPTG and supplemented with 1.5% glucose. **(c)** Gluconate titer as a function of time. Error bars represent 1 s.d. from the mean of three technical replicates (n=3). **(d)** Gluconate titer following 18.25 hr of induction.

**Figure 3.**
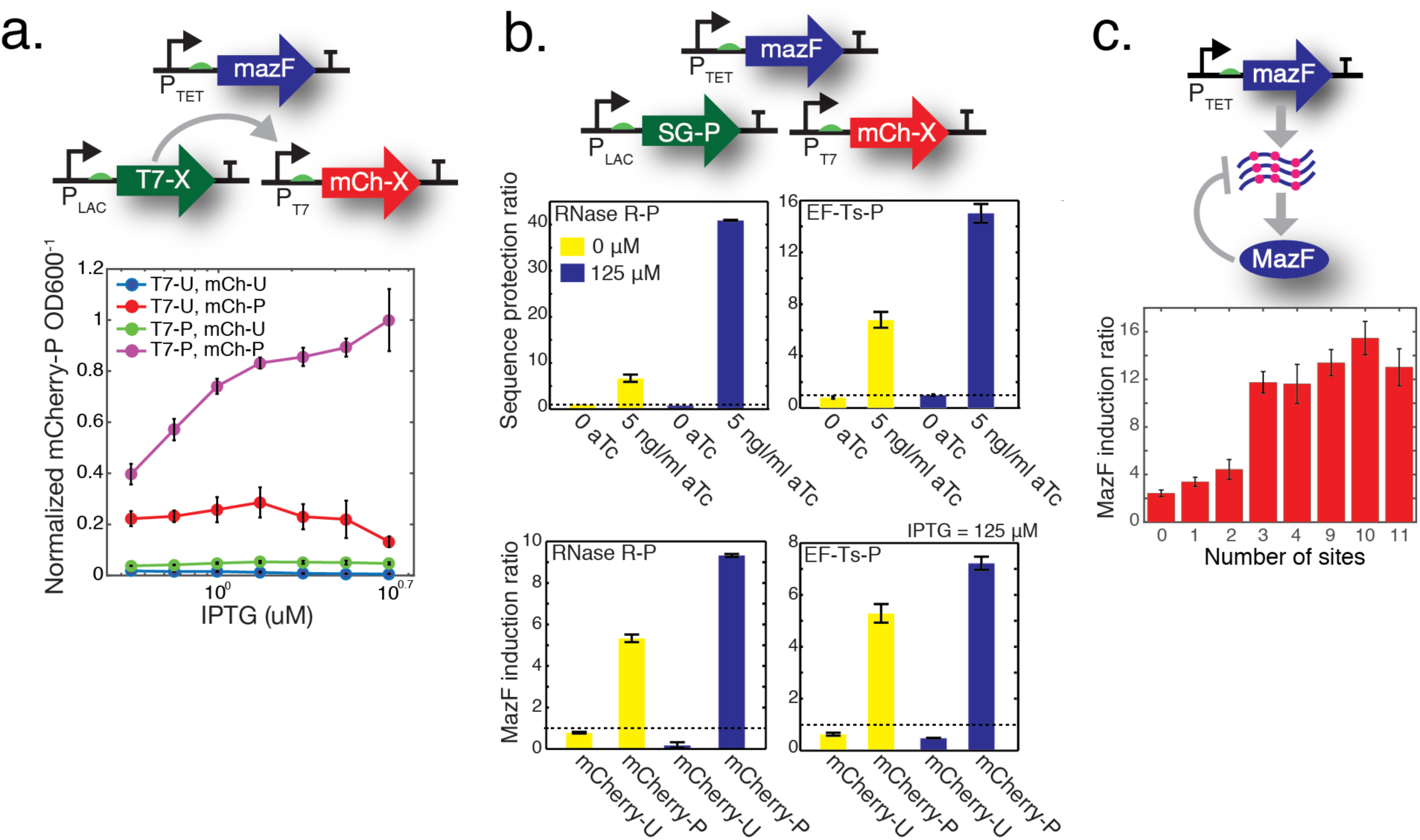
Optimization of the MazF resource allocator performance by protection of support genes and negative feedback loop strength. **(a)** Schematic of T7 polymerase circuit (top). MazF, T7-X and mCherry-X were controlled by an aTc (P_TET_), IPTG (P_LAC_) or T7 (P_T7_) regulated promoter. Unprotected T7 (T7-U) contained 50 sites. Normalized fluorescence as a function of IPTG for cells expressing all combinations of T7-U, T7-P, mCherry-U and mCherry-P following 8.3 hours of induction (bottom). Error bars represent 1 s.d. (n = 3). **(b)** Schematic of support gene circuit (top). MazF, protected support gene (SG-P) and mCherry-X were controlled by P_TET_, P_LAC_ or P_BAD_. Sequence protection ratio (middle) for cells in the presence or absence of IPTG or aTc. Cells were induced for 8.3 hr. MazF induction ratio (bottom) in the presence (5 ng/ml aTc, 125 uM IPTG) or absence (0 ng/ml aTc, 0 ng/ml IPTG) of IPTG or aTc. Cells were induced with 0.05% arabinose for 8.3 hr. Error bars represent 1 s.d. (n = 4). **(c)** Schematic of MazF mRNA-decay feedback loop (top). MazF induction ratio for cells expressing *mazF* transcripts that varied in the number of recognition sites (P37-43 in Supplementary Table I). Cells were induced with 0 or 5 ng/ml aTc and 0.05% arabinose for 9.2 hr. Error bars represent 1 s.d. (n = 4).

To measure if the expression of mCherry-P degraded as a function of time in MazF-induced cells, cell populations were induced with mCherry-P at three time points following exposure to MazF. To compare expression across conditions, mCherry-P fluorescence was divided by biomass and normalized to the maximum expression level across all conditions following 12 hr of induction with 5 ng/ml aTc. In the absence of MazF, delayed induction by 2 hr reduced mCherry-P expression by 85% (Supplementary Figure 4a), whereas cells induced with MazF displayed a 34% decrease in mCherry-P expression (Supplementary Figure 4b). These data indicate that heterologous expression was significantly attenuated by delayed induction in the absence of MazF. By contrast, delays in the induction of mCherry-P reduced expression by a smaller magnitude in the presence of MazF, indicating that MazF-induced cells preserved high metabolic activity for a period of time.

**Figure 4.**
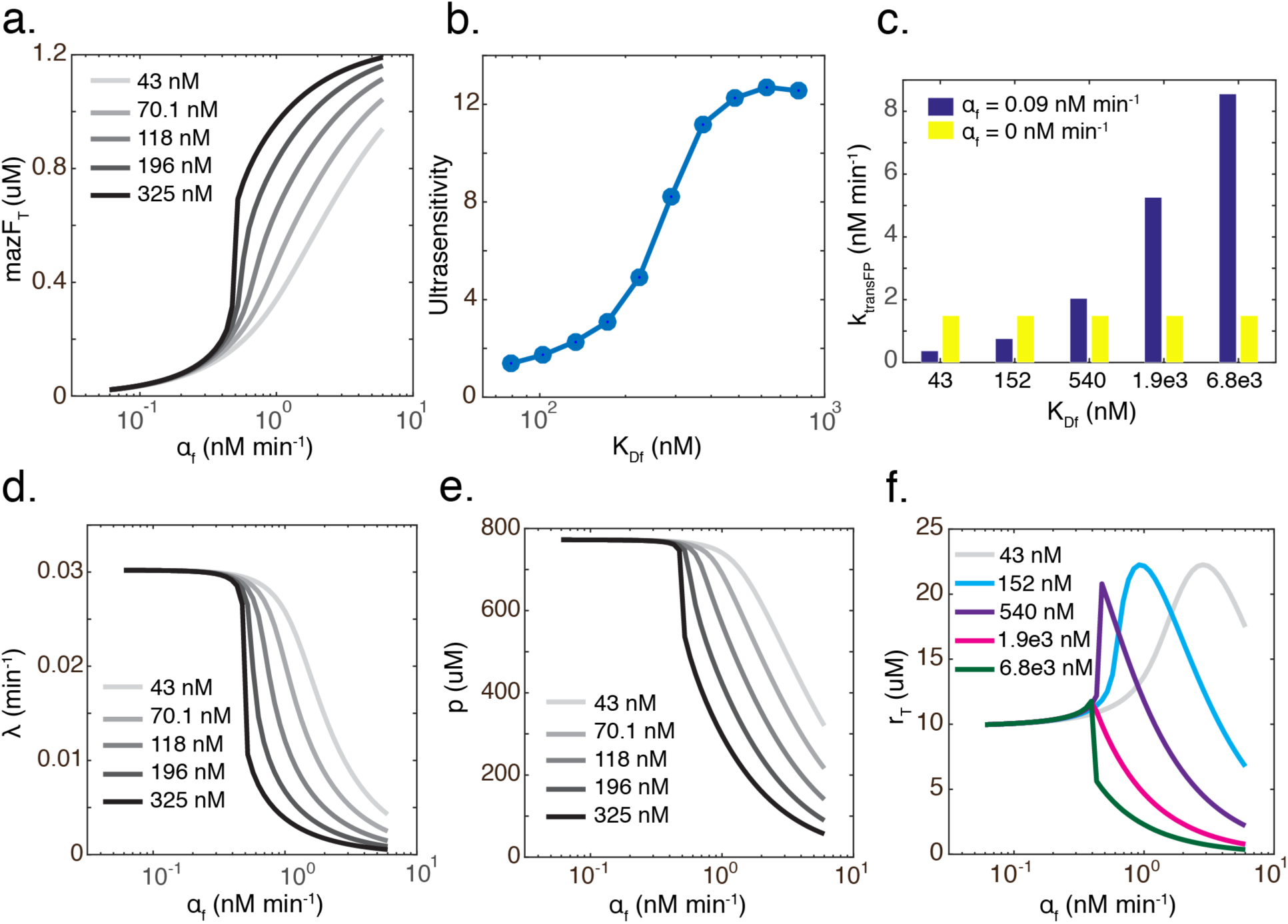
A dynamic resource allocation model demonstrates that the MazF mRNA-decay feedback loop established proportional control of the concentration of steady-state MazF. **(a)** Total MazF concentration at steady-state (mazF_T_, t = 278 hr) as a function of the transcription rate of *mazF* (α_f_) across a range of dissociation constants (K_Df_) of MazF to *mazF* mRNA (m_f_). **(b)** Maximum logarithmic sensitivity (ultrasensitivity) of the dose response of α_f_ vs. mazF_T_ across a range of K_Df_ values. **(c)** Steady-state translation rate of a protected reporter gene FP (k_transFP_) as a function of K_Df_ in the presence (α_f_ = 0.09 nM min^−1^) or absence (α_f_ = 0 nM min^−1^) of MazF. **(d)** Steady-state growth rate () as a function of α_f_ for different values of K_Df_. **(e)** Steady-state unprotected proteome (*p*) concentration as a function of α_f_ for different values of K_Df_. **(f)** Steady-state total ribosome concentration (r_T_) as a function of α_f_ for different values of K_Df_.

To distinguish whether transcriptional or translation activity dominated the enhancement of mCherry-P in response to MazF, *mCherry-P* mRNA was measured using quantitative real-time PCR (qPCR). The *mCherry-P* mRNA fold change following 56 min of induction with 0 or 5 ng/ml aTc relative to *mCherry-P* mRNA abundance at the beginning of the experiment (t = 0) was similar in the presence or absence of MazF (Supplementary Figure 5). These data show that MazF did not significantly alter the *mCherry-P* transcription rate over this period of time. Therefore, these results suggest that MazF activity augmented the translation rate of *mCherry-P* relative to *mCherry-U*.

**Figure 5.**
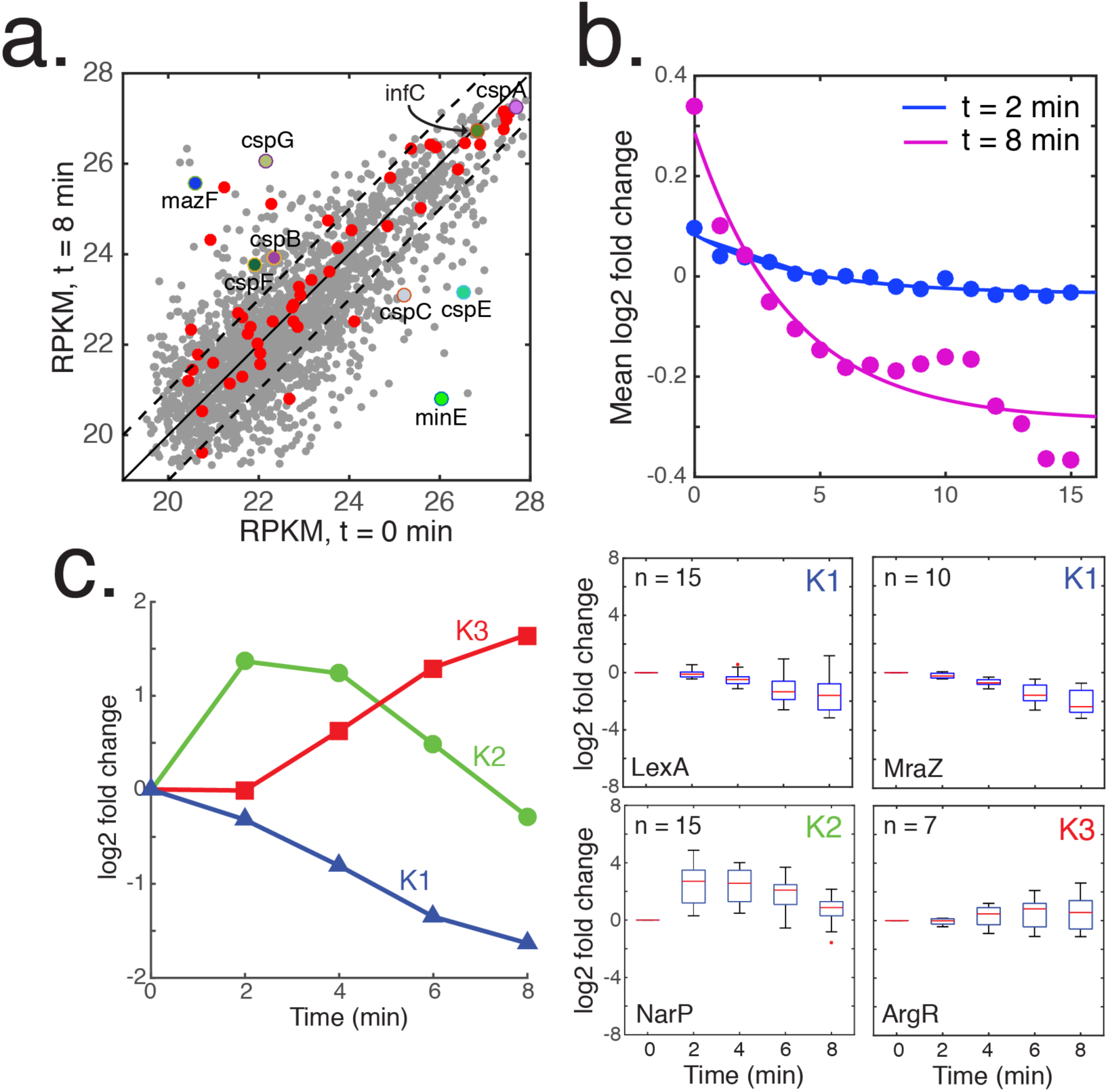
Time-series RNA-seq measurements of MazF-induced cells identified the relationship between transcript abundance and the number of MazF recognition sites and provided insight into physiological impact of MazF expression. **(a)** Scatter plot of RPKM measurements prior to induction with MazF vs. induction with MazF (5 ng/ml aTc) for 8 min. Gray and red data points denote unprotected or protected transcripts larger than 80 nucleotides. Dashed lines represent a two-fold threshold in transcript abundance. *cspABCGEF*, *mazF* and *minE* transcripts are highlighted. **(b)** Scatter plot of the number of *mazF* sites for each gene vs. mean log2 fold change following induction with 5 ng/ml aTc for 2 or 8 min. A 5-point moving average was applied to the data. Lines represent fitted exponential functions to the data. **(c)** K-means clustering of log2 fold change of genes (left) that exhibited correlated dynamics between biological replicates (n = 939). Box plots (right) of representative functional or regulatory enrichments in the K1, K2 and K3 clusters according to the Fisher's exact test (p < 0.05). On each box, the red line indicates the median, the bottom and top edges represent the 25^th^ and 75^th^ percentiles and ‘+’ denote outlier data points.

### Enhancement of gluconate production using the MazF resource allocator

The gluconate pathway competes directly with biomass synthesis by redirecting glucose into gluconate via glucose dehydrogenase (Gdh, Figure 2a). To determine the impact of MazF on metabolic flux, biomass and gluconate were measured as a function of time (see Materials and Methods) in cells expressing protected Gdh (*gdh-P*) or unprotected Gdh containing 10 sites (*gdh-U*) controlled by a P_LAC_ promoter. These experiments were conducted in a strain background that contained genetic modifications to inhibit gluconate metabolism and decouple glucose phosphorylation and transport to efficiently utilize glucose as a substrate for target metabolic pathways (KTS022IG, mazF::Δ, see Materials and Methods)^15^.

Cell growth was inhibited by MazF induction whereas the uninduced population continued to grow as a function of time (Figure 2b). Cells bearing *gdh-P* driven by a P_LAC_ promoter displayed up to a 3-fold higher gluconate concentration and 5-fold higher gluconate per unit time in the presence of MazF compared to cells that were not induced with aTc (Figure 2c and Supplementary Figure 6a). The gluconate titer was 85% higher for cells induced with MazF and Gdh-P compared to cells that were not induced with aTc following 18.25 hr (Figure 2d). A protected fluorescent reporter (sfGFP-P) N-terminally fused to Gdh-U or Gdh-P increased up to 3.3 and 5-fold as a function of aTc (Supplementary Fig. 6b). These data demonstrated that the MazF resource allocator could enhance metabolic flux by protecting genes in a target metabolic pathway.

### Enhancement of resource redistribution activity by protection of key host-genes that support synthetic circuit operation

Synthetic circuits depend on a dense network of host-genes including the transcriptional and translational machinery. Therefore, MazF-mediated decay of host factors could lead to degradation of circuit function. To investigate whether protection of support genes could improve the performance of the resource allocator, we tested whether protection of an orthogonal RNA polymerase T7 could enhance the circuit output. A protected (T7-P) or unprotected (T7-U, 50 sites) T7 controlled by an IPTG-inducible promoter (P_LAC_) was used to drive the expression of mCherry (Figure 3a). The combination of T7-P and mCherry-P yielded a 21 or 7.6-fold higher expression level of mCherry compared T7-P, mCherry-U or T7-U, mCherry-P in the presence of MazF (5 ng/ml aTc) and 1 mM IPTG. T7-P regulating an N-terminal fluorescent protein fusion of mCherry-P to Gdh-P (mCherry-P-Gdh-P) exhibited a 1.4 and 15-fold higher expression compared to T7-P, mCherry-P-Gdh-U or T7-U, mCherry-P-Gdh-P (Supplementary Figure 7). The mCherry expression level of the T7-X, mCherry-X (Figure 3a) and T7-X, mCherry-X-Gdh-X (Supplementary Figure 7) circuits were differentially enhanced by protection of T7 or the reporter gene (*mCherry-X* or *mCherry-X-gdh-X*) in the presence of MazF.

Thus, the enhancement of resource redistribution activity by protection of specific genes in a circuit depended on the circuit composition.

Identifying translation factors in need of protection is challenging since the basic translation machinery consists of 78 factors including ribosomal proteins and aminoacyl-tRNA synthases^16^. To identify candidates, the proteome of MazF-induced cells was measured as a function of time. The majority of the proteome (216 measured proteins) and 91% of 35 detectable ribosomal proteins varied by less than 10% following 5 hours of induction, demonstrating that highly abundant proteins were stable following exposure to MazF (Supplementary Figure 8a). Ribosomal protein subunits S9, S20 and L17 decreased by ~20% and an essential elongation factor EF-Ts decreased by approximately 80% following 5 hours of induction with MazF (Supplementary Figure 8b). In the presence of MazF, a protected version of EF-Ts (EF-Ts-P) driven by an IPTG-dependent promoter (P_LAC_) significantly enhanced the expression of mCherry-P compared to cells that were not induced with EF-Ts-P (Figure 3b). These results indicated that genome-wide measurements could be used to discover support genes in need of protection to augment resource redistribution activity.

Global mRNA decay could generate imbalances in the expression levels of genes in a regulatory network. For example, high concentrations of truncated mRNA fragments could saturate exonucleases that process these fragments into mononucleotides^17^. Further, mRNA cleavage generates ribosome stalling at the 3’ end of the mRNA, referred to as non-stop complexes, which require the action of ribosome recycling factors to rescue the ribosomes^18^. RNase R is a multifunctional protein that exhibits ribonuclease and ribosome recycling factor activities^19^. Co-expression of MazF and RNase R-P significantly enhanced the expression of mCherry-P compared to cells expressing only MazF (Figure 3b). However, co-expression of EF-Ts-P and RNase R-P did yield an additional enhancement in the level of mCherry-P in the presence of MazF compared to cells expressing either of single support genes, RNase R or EF-Ts-P (Supplementary Figure 9). These results suggested that epistasis among support genes could potentially limit incremental improvement of resource redistribution activity.

### The mazF mRNA-decay feedback loop enhanced resource redistribution activity via proportional control

The *mazF* transcript is enriched for recognition sites (Supplementary Figure 1b), establishing an mRNA-decay negative feedback loop. We suspected that protection of MazF could enhance circuit performance. However, the feedback loop may modulate the regulatory dynamics of MazF and therefore influence resource redistribution activity. To investigate this possibility, we probed the role of the feedback in the MazF resource allocator.

Cells induced with *mazF-U* (9 sites) exhibited a lower steady-state mRNA level compared to cells expressing *mazF-P* (Supplementary Figure 10a), demonstrating that the feedback loop was actively regulating the abundance of the *mazF* transcript. Corroborating this result, a 35% lower threshold of aTc was required to inhibit growth in a strain expressing MazF-P compared to MazF-U (Supplementary Figure 10b), suggesting that protection of *mazF* mRNA enhanced the MazF protein level. The Hill coefficients of OD600 as a function of aTc following 11.2 hr of induction were 2.6 and 5.9 for cells induced with MazF-U or MazF-P, revealing an ultrasensitive relationship between MazF activity and cell growth.

Contrary to expectation, MazF-U displayed significantly higher mCherry-P expression compared to MazF-P across a broad range of aTc concentrations, showing that the negative feedback loop enhanced resource redistribution activity (Supplementary Figure 10c). To further investigate the quantitative relationship between feedback loop strength and resource redistribution activity, we varied the number of recognition sites in the *mazF* transcript (Figure 3c, Supplementary Figure 11). The MazF induction ratio increased with the number of sites and the wild-type *mazF* sequence (9 sites) generated nearly the highest output level. In sum, these results indicated that the activity of the feedback loop was a tunable knob that could be used to modulate circuit performance.

A dynamic resource allocation model was constructed to provide insight into the role of the negative feedback loop on circuit behavior (Supplementary Text). The model represented the mRNA and protein levels of key species, which compete for limiting ribosome pools including ribosomes (*r*), unprotected proteome (*p*), MazF (*mfp*) and a protected reporter gene (*FP*). The growth rate () function was based on a previous coarse-grained mechanistic model of gene expression and growth^20^. The model equations are described in the Supplementary Text and Supplementary Table II and III provides a list of model species and parameters.

The relationship between steady-state total MazF concentration (mazF_T_) and the FP translation rate (k_transFP_ = k_trans_FP) is non-monotonic (Supplementary Figure 12a), indicating that there is an optimal level of mazF_T_ to maximize the synthetic circuit output. The model shows that the strength of the feedback loop, represented by the dissociation constant of MazF dimer (*mfpd*) to the *mazF* transcript *mf* (K_df_ = k_rff_/k_f_), is inversely correlated with the dose-response ultrasensitivity of mazF_T_ as a function of the *mfp* transcription rate (α_f_, Figure 4a,b). Molecular mechanisms that realize ultrasensitivity include MazF dimerization^21^, molecular sequestration^22^,^23^ of mRNAs by ribosomes^24^, or positive feedback^25^. In addition, thresholded control of  by mazF_T_, which was observed in our experimental and modeling data (Figure 4d and Supplementary Figure 10b), could contribute to ultrasensitivity in the network. For high K_Df_ corresponding to the open loop system, the model exhibits bistability manifesting as two stable steady states across a range of *α_f_* values (Supplementary Figure 12b). These results show that the negative feedback loop enables proportional adjustment of the mazF_T_^26^ and reduces the potential for bistability by abolishing ultrasensitivity^23^,^27^ (Figure 4b). As such, mazF_T_ could be tuned to operate in the regime that maximized resource redistribution activity.

For a fixed value of α_f_, k_transFP_ is inversely related to K_Df_ (Figure 5f), qualitatively recapitulating the increase in mCherry-P with the number of binding sites in the *mazF* transcript (Figure 3e).  and the concentration of *p* decrease as a function of α_f_, mirroring experimental data that showed a decrease in saturating OD600 and mCherry-U as a function of aTc (Supplementary Figure 2, 3a, 10b and Figure 4c,d). The increase in ultrasensitivity of the dose response of mazF_T_ vs.  with the strength of the feedback loop (Figure 4d) reflected the enhanced ultrasensitivity of the steady-state dose response of aTc vs. biomass (OD600) for cells expressing MazF-P compared to MazF-U (Supplementary Figure 10b). Reducing the negative feedback loop activity narrows the range of α_f_ values that map to high total *r* concentration (r_T_). Above a threshold value of K_Df_ (K_Df_ ≥ 1.9 = M), r_T_ decreases monotonically with α_f_ (Figure 4e). The negative feedback has important implications for resource allocator design by enabling precise tuning of the MazF operating point by establishing a proportional relationship between α_f_ and mazF_T_. Indeed, this negative feedback may provide an evolutionary advantage for cells by preventing the deleterious effects of MazF overexpression that accelerated cell death (Supplementary Figure 13).

### Time-series RNA-seq measurements provided insights into the physiological impact of MazF activity and identified design elements for building the MazF resource allocator

To provide insights into the genome-wide variation in transcript abundance following MazF exposure, RNA-seq measurements were collected every 2 min for a total of 8 min. The majority of the 192 endogenous protected genes increased or remained constant following induction with MazF for 8 min (Figure 5a). The abundance of each transcript is determined by a balance between the rates of synthesis and decay catalyzed by RNases and MazF. Therefore, it was challenging to decipher the direct effects of MazF on transcript abundance. Nevertheless, the number of MazF sites was negatively correlated with the mean log2 fold change of transcript abundance following 8 min of induction with aTc, indicating that on average the number of MazF sites predicted the fold-change across the transcriptome (Figure 5b).

Partitioning the transcriptome fold-change dynamics into three clusters (see Materials and Methods) identified transcripts that were down-regulated (K1, n = 460), rapidly increased and delayed down-regulation (K2, n = 148) or increased in response to MazF induction (K3, n = 331, Figure 5c and Supplementary Figure 14). We evaluated functional or regulatory enrichments (p < 0.05) in each cluster to provide insights into the physiological response to MazF exposure (Supplementary Table V). Cell envelope and genes regulated by Fur, MraZ and LexA were enriched in the K1 cluster (Figure 5c and Supplementary Figure 14b). MraZ is a transcriptional repressor that controls many genes involved in cell division and cell wall biosynthesis^28^. In addition, the cell division regulator *minE* mRNA decreased significantly in the RNA-seq data (Figure 5a), corroborating a link between MazF activity and inhibition of cell division^29^,^30^. The K2 cluster was enriched for genes regulated by NikR, GlpR, GcvA, IHF, IscR and RstA and amino acid and anaerobic metabolism (Supplementary Figure 14). The numerous regulatory categories in K2 (Supplementary Table V) suggested that the pulsatile transcript dynamics could be established by an initial increase in synthesis due to changes in transcription factor activity and delayed down-regulation due to mRNA-decay at a threshold concentration of MazF. Genes regulated by ArgR were enriched in the up-regulated cluster K3. In addition, 11 TCA cycle enzymes were up-regulated in the RNA-seq data (p = 0.051 enrichment in K3), suggesting that MazF-induced cells exhibited high metabolic activity (Supplementary Figure 15 and Supplementary Table V). Previous work has demonstrated that fumarate production increased the frequency of persister cells following antibiotic exposure^31^. A closer examination of the catabolic pathway revealed that fumarate producing enzymes were significantly induced, suggesting that MazF activity could be connected to persistence via enhancement of fumarate flux^32^,^33^ (Supplementary Figure 15).

Cold-shock genes are selectively expressed in response to cold stress and perform diverse functions including unwinding of RNA secondary structures, modulation of ribosome and DNA/RNA chaperone activity^34^. The RNA-seq data demonstrated significant shifts in cold-shock *cspBCEFG* and associated *rbfA*, *rhlB*, *rhlE* and *deaD* transcript abundance as a function of time (Supplementary Figure 16). IF-3, one of the major translation factors in *E. coli*, has been shown to mediate cold shock translational bias in response to cold stress^35^,^36^. IF-3 increased over 4-fold in the proteomics data (Supplementary Figure 8b) following 5 hr of MazF induction, whereas the abundance of *infC* mRNA did not change significantly in the RNA-seq data (Figure 5a and Supplementary Figure 16). Future work should investigate the molecular mechanisms that underlie the connection among MazF activity, up-regulation of IF-3, and rapid changes in cold-shock mRNA abundance.

Since cold-shock transcripts were up-regulated in response to MazF activity, these sequences were promising candidates for engineering MazF-responsive promoters. To test the modularity of cold-shock induction by MazF, we constructed a tandem promoter composed of P_LAC_ upstream of the *cspB* or *cspG* promoter, UTR and the first 14 amino acids of CspG or CspB N-terminally fused to sfGFP-P (Supplementary Figure 17). MazF induction increased sfGFP-P by a maximum of 20 or 80-fold, demonstrating that the *cspB* and *cspG* regulatory sequences are modular control elements that directly respond to MazF activity as an input.

### Relationship between MazF cleavage efficiency and predicted mCherry mRNA secondary structure

A quantitative understanding of the mapping between MazF site placement and cleavage efficiency could enable tuning of the timing and degrees of protection to inform resource allocator design. To explore the dominant parameters that influence MazF cleavage efficiency, we varied the number and position of MazF recognition sites in the *mCherry* transcript. Previous work demonstrated that MazF activity was inhibited by strong secondary structures and ribosomes enhanced cleavage efficiency by unwinding mRNA secondary structures during translation^37^.

To map the relationship between position and cleavage efficiency, a single MazF site was inserted at 14 positions in *mCherry-P* (Supplementary Figure 18). These *mCherry* sequences exhibited a broad range of expression levels in response to MazF (Supplementary Figure 19a). The output was correlated with the predicted secondary structure Gibbs free energy (∆G) 40-47 bp upstream of the recognition site (ρ ranged between −0.7 to −0.6, p < 0.01) computed using NUPACK (Supplementary Figure 18b,c). For sequences spanning upstream and downstream of the MazF site, mCherry expression was correlated (ρ = −0.6, p < 0.05) with ∆G (38-41 bp, Supplementary Fig. 18d). However, the ∆G of the sequence downstream of the recognition site was not correlated with the expression level of mCherry across a broad range of window sizes (Supplementary Fig. 18e). Therefore, MazF cleavage efficiency could be predicted using the folding energy of the local mRNA secondary structure upstream or across the recognition site.

To provide insight into the programmability of MazF cleavage efficiency, we interrogated whether measurements of *mCherry* variants containing a single MazF site (Supplementary Figure 18a) could predict the expression of *mCherry* sequences containing combinations of sites. mCherry expression decreased as a function of the number of recognition sites in the presence of MazF (Supplementary Fig. 18f). The product of the single site *mCherry* expression levels could predict the expression of the multi-site variants (*p* < 0.0001), suggesting that combinations of MazF recognition sites could be used to modulate the degree of transcript protection.

## DISCUSSION

A major goal of synthetic biology and metabolic engineering is to develop strategies to control the resource economy of cells for switching between modes of growth and production^38^. During a production phase, cellular energy and resources are focused on specific pathways, while minimizing resource expenditure towards nonessential cellular operations. Towards these objectives, previous work leveraged tunable enzymatic degradation of a metabolic hub that determines the direction of metabolic flux to augment the yield and titer of a metabolic pathway two-fold^39^. While this strategy provided localized control of metabolic flux, it does not modulate the global allocation of subsystems such as transcription and translation. On a larger scale, inducible regulation of RNA polymerase subunits was recently used to control *E. coli* growth^36^. However, this mechanism cannot be generally applied to redirect resources towards engineered networks. Here, we achieved programmable and genome-wide manipulation of intracellular resources to enhance a target function by exploiting global mRNA decay. This approach could be harnessed for diverse applications by systematically discovering genes in need of protection to enhance a target function. Indeed, recent advancements in DNA synthesis technologies will facilitate large-scale recoding of support genes to protect from MazF activity.

MazF regulates orders of magnitude more genes simultaneously compared to other technologies such as CRISPRi. Therefore, this approach could be used to reprogram cellular behavior in response to changing environments^41^,^42^. Homologues of MazF that recognize 3, 5 and 7-bp recognition sites have been identified in diverse bacterial species^43–45^. Active site mutations have been shown to modify the MazF sequence specificity, suggesting that protein engineering could be used to expand the diversity of MazF recognition site sequences^46^. The variation in recognition sequence specificity could be used to tune the number of genes targeted by MazF.

In the model, the MazF transcription rate *α_f_* is a bifurcation parameter that triggers bistability in the absence of negative feedback (Supplementary Figure 19b). Bistability can be established in circuit with ultrasensitivity and positive feedback^23^. Positive feedback can originate from several mechanisms, which increases the MazF translational rate. For *α_f_* = 0, increasing the transcription rate of *p* (*α_p_*) reduces r_T_, which reveals that *mp* and *mr* compete for limiting ribosome pools (Supplementary Figure 19c). Since MazF has a 2-fold higher binding affinity for *mp* compared to *mr*, reducing *mp* levels yields an increase in the translation rate of *r* and hence the translation rate of MazF, forming a net positive feedback loop. The strength of the positive feedback loop could also be modulated by changes in protein concentrations due to growth rate inhibition. The *mazF* mRNA-decay negative feedback realizes a proportional relationship between *α_f_* and the total concentration of MazF at steady-state, thus reducing the potential for bistability in the network^23,26,27^. Future work should explore how negative feedback established by mRNA-decay modulates the sensitivity of a circuit to parameter variations and response time compared to transcriptional autoregulation^47^,^48^.

There are several challenges and limitations to optimizing the MazF resource allocator. MazF activity increased the abundance of a set of host-cell transcripts, which sequesters resources away from engineered circuits. However, this activation program could be exploited by repurposing regulatory elements that respond to MazF activity to expand the resource allocator design. In addition, MazF activity has been shown to yield a heterogeneous ribosome pool by targeting a specific site of the 16S rRNA^49^, which could manifest as translation bias for selected transcripts^50^. Decay of the unprotected proteome occurs on the timescale of hours, thus limiting the time scale required to shift metabolic flux. To rapidly manipulate metabolic flux, induction of MazF could be coupled with proteases^7^ for targeted control of protein abundance. As the proteome decays, stoichiometric relationships required for protein activity must be maintained^51^. Further, MazF has been shown to establish a futile cycle of continuous RNA synthesis and decay, resulting in energy dissipation^32^. To minimize an energy deficit, orthogonal T7-P could be used to drive the engineered pathway while inhibiting native RNA polymerases. Future efforts should pinpoint the dominant parameters that influence resource redistribution activity.

Top-down approaches such as MazF could be used to discover host factors that preserve high metabolic activity in the absence of growth. Genome engineering could be used to protect these pathways from MazF activity^52^. Optimal regulatory strategies should be designed to balance enhancement of resource redistribution activity and degradation of cellular support subsystems over long time scales. For example, MazF could be transiently induced until energy degrades to a threshold that triggers inhibition of MazF activity and allows rebalancing of the proteome^53^. Together, advances in regulatory control strategies and large-scale recoding could enable the design of protected and unprotected orthogonal sub-genomes that dynamically switch between cellular operations.

## MATERIALS AND METHODS

### Cloning and strain construction

*mazF* was deleted from the *E. coli* BW25113 strain using lambda-red recombination and verified by colony PCR. MazF was introduced into an intergenic region referred to as SafeSite 1 (chromosomal position 34715) under control of an aTc-inducible promoter (P_TET_). PCR amplifications were performed using Phusion High-Fidelity DNA Polymerase (NEB) and oligonucleotides for cloning and strain construction were obtained from Integrated DNA Technologies. Standard cloning methods were used to construct plasmids. Plasmids were based on a previously generated construct library^54^. A list of plasmids and strains used in this study can be found in Supplementary Table I.

### Growth conditions and plate reader experiments

For plate reader experiments, cells were grown at 37°C for approximately 6-8 hours and then diluted to OD600 of 0.01 in a 96-well plate (Corning) in LB Lennox media (Sigma). For plate reader experiments, cells were grown in 200 µl volumes at 37°C in 96-well plate covered by a gas-permeable seal (Fisher Scientific) in an M1000 (Tecan) or Synergy 2 (BioTek) plate reader. The method measured cell density (OD600) and fluorescence every 10 min for 15 hr. The M1000 excitation and emission wavelengths were 485, 510 nm for GFP and 587, 610 nm for RFP (5 nm bandwidth). The BioTek excitation and emission wavelengths were 485, 528 nm for GFP and 560, 620 for RFP (20 nm bandwidth). The M1000 and Synergy 2 measured absorbance at 600 nm (OD600) to quantify total biomass.

### qPCR measurements

Oligonucleotides for quantitative real-time PCR (sequences are listed in Supplementary Table IV) were designed using Integrated DNA Technologies. 500 ng of total RNA was reverse transcribed using the iScript cDNA synthesis kit (Bio-Rad). The reaction mix contained 5 µl of SsoAdvanced Universal Probes Supermix (Bio-Rad), 0.5 µl primer and probe corresponding to 250 nM primers and 125 nM probe (20X stock) and 0.5 µL of cDNA. qPCR measurements were performed using a CFX96 real-time PCR machine (Bio-Rad). The relative expression levels were determined by a 2^−ΔΔG^ method. Each sample was normalized by the cycle threshold geometric mean using reference genes *rrsA* and *cysG*^55^.

### Gluconate measurements

KTS022IG mazF::Δ (strain S1 in Supplementary Table I) strains bearing pBbA6c-gdh-X (plasmid P8-9 in Supplementary Table I) and pBbS2k-mazF-U (plasmid P1) were grown in LB medium at 37°C overnight and used to inoculate a 10 mL culture the next morning at an OD600 of 0.05. At OD600 of 0.3, 1.5% glucose, 1000 uM IPTG and 5 or 0 ng/ml were administered to the cultures. 1 mL samples were collected at the specified times and centrifuged at 5000 × g for 5 min to isolate the supernatant. The supernatant samples were analyzed for gluconic acid using a 1200 Series liquid chromatography system (Agilent Technologies, Santa Clara, CA) coupled to an LTQ-XL ion trap mass spectrometer (Thermo Scientific, San Jose, CA) equipped with an electrospray ionization source. Aliquots of the diluted samples were injected onto a Rezex ROA-Organic Acid H+ (8%) (150 mm × 4.6 mm) column (Phenomenex, Torrance, CA) equipped with a Carbo-H+ (4 mm × 3 mm) guard column (Phenomenex). Gluconic acid was eluted at 55 °C at approximately 3.5 min with an isocratic flow rate of 0.3 mL/min of 0.5% (v/v) formic acid in water. Precursor ion m/z 195.1 was selected in negative ion mode using an isolation window of m/z 2 and was fragmented with a normalized collision energy of 35. Fragment ions were analyzed in the range of m/z 50-200. m/z 128.5-129.5 was selected as pseudo-MRM transition for compound quantification. Resulting peak areas were compared to an external standard calibration in the range of 0.2-200 uM. The source parameters were ion spray voltage: 4 kV; capillary temperature: 350 °C; capillary voltage: −2 V; tube lense voltage: −40 V; sheath gas flow: 60; auxiliary gas flow: 5; and sweep gas flow: 10 (all arbitrary units).

### Proteomics

BW25113 mazF::Δ, SafeSite1::tetR-P_TET_-mazF (strain S2 in Supplementary Table I) was grown overnight in LB at 37°C and then diluted to an OD600 of 0.05 in a 500 ml LB culture. At OD600 of 0.5, cell populations were induced with 5 ng/mL aTc and 40 mL of cells were collected approximately every hour and centrifuged for 5 min at 4300 × g. Proteomic samples were prepared for analysis as previously described^56^. Briefly, the cell pellets were lysed and proteins were extracted by chloroform/methanol precipitation. The proteins were resuspended in 100 mM AMBIC with 20% methanol and reduced with tris(2-carboxyethyl)phosphine (TCEP) for 30 min, followed by addition of iodoacetamide (IAA; final conc. 10 mM) for 30 min in the dark, and then digested overnight with MS-grade trypsin (1:50 w/w trypsin: protein) at 37°C. Peptides were stored at −20°C until analysis.

Samples were analyzed on an Agilent 1290 UHPLC - 6550 QTOF liquid chromatography mass spectrometer (LC-MS/MS; Agilent Technologies) system and the operating parameters for the LC-MS/MS system were described previously^56^. Peptides were separated on a Sigma-Aldrich Ascentis Express Peptide ES-C18 column (2.1 mm × 100 mm, 2.7 mm particle size, operated at 60°C) and a flow rate of 0.4 mL/min. The chromatography gradient conditions were as follows: from the initial starting condition (98% buffer A containing 100% water, 0.1% formic acid and 2% buffer B composed of 100% acetonitrile, 0.1% formic acid) the buffer B composition was held for 2 min then increased to 10% over 3 min; then buffer B was increased to 40% over 117 min, then increased to 90% B over 3 min and held for 8 min, followed by a ramp back down to 2% B over 1 min where it was held for 6 min to re-equilibrate the column to the original conditions. The data was analyzed with the Mascot search engine version 2.3.02 (Matrix Science) and filtered and validated by using Scaffold v4.3.0 (Proteome Software Inc.) as described previously^56^.

### RNA-seq library construction and sequencing

BW25113 mazF::Δ, SafeSite1::tetR-P_TET_-mazF (strain S2 in Supplementary Table I) was grown overnight in LB at 37°C and then diluted to an OD600 of 0.05 in a 10 mL LB culture. At an OD600 of 0.5, cells were induced with 5 ng/mL aTc. Samples were collected as follows: 200 µl of the cell cultures were added to 400 µl of RNAprotect (Qiagen) to stabilize the RNA, incubated for 5 min at room temperature and then spun down for 10 min at 5000 × g. Total RNA was isolated using RNeasy purification kit and treated with DNAase I (Qiagen). The Functional Genomics Lab (FGL), a QB3-Berkeley Core Research Facility at UC Berkeley, constructed the sequencing libraries. At the FGL, Ribo-Zero rRNA Removal Kits (Illumina) were used to remove ribosomal RNA and ERCC RNA Spike-In Control Mixes (Ambion by Life Technologies) were added to the samples. The library preparation was performed on an Apollo 324™ with PrepX™ RNAseq Library Prep Kits (WaferGen Biosystems, Fremont, CA), and 18 cycles of PCR amplification was used for index addition and library fragment enrichment.

### RNA-seq data analysis

The read counts were mapped onto the MG1655 genome using Bowtie 1^57^ on the galaxy webserver^58^. Reads per kilobase of transcript per million (RPKM) was computed by multiplying the number of mapped reads by 10^9^ and then dividing by the gene length and median number of total reads for each condition. For clustering analysis, the correlation coefficient (ρ = 0.9) between two biological replicates as a function of time was used as a threshold to remove genes that exhibited variability between replicates. The log2 fold change was partitioned into clusters using the K-means algorithm (MATLAB). To determine an optimal number of clusters, the sum of squared errors (SSE) was computed for each data point from the corresponding cluster centroid across a range of K-values (1-10). The Elbow method was used as a heuristic to select the optimal number of partitions that minimizes the SSE. The Fisher’s exact test (p < 0.05) was used to evaluate enrichment of genes based on TIGRFAM annotation (MicrobesOnline) or transcription factor network (RegulonDB). Supplementary Table V contains a list of genes in the enriched categories.

### Computational modeling

We used custom code for computational modeling and data analysis in MATLAB (Mathworks) and Python. Details about the model construction are provided in the Supplementary Text. Model species and parameters are listed in Supplementary Tables II and III.

### Characterization of cell viability

BW25113 mazF::Δ strains transformed with pBbS2k-mazF-U or pBbS2k-mazF-P were grown overnight at 37°C in LB media and then diluted to an OD600 of 0.01 in 5 ml LB media. At an OD600 of 0.3, 5 ng/mL aTc dissolved in 100% ethanol was used to induce *mazF* and an equivalent volume of 100% ethanol was administered to the uninduced cell populations. Following 0 and 7 hr, cells were prepared for fluorescent microscopy using the LIVE/DEAD Baclight™ Bacteria Viability Kit (Thermo Fisher) to characterize the fraction of viable cells across the population as a function of time. Microscope images were collected using a Zeiss Axio Observer D1 and Plan-Apochromat 63/1.4 Oil Ph3 M27 objective. Cells were imaged using excitation BP 470/40 and emission BP 525/50 (Filter Set 38 High Efficiency) or excitation 560/40 and emission BP 630/75 (Filter Set 45). Images were captured with a Hamamatsu ORCA-Flash4.0 using the ZEN Software (Zeiss). Cell Counter (Fiji)^59^ was used to analyze the images and quantify the number of viable and dead cells.

## ACKNOWLEDGEMENTS

We would like to thank Karen Lundy for constructing the RNA-seq libraries and Kristala Prather (MIT) for providing *E. coli* strain S1 (see Supplementary Table I) for the gluconate measurements. This work was supported by the US Department of Energy (Grant DE-SC0008812) and used the Vincent J. Coates Genomics Sequencing Laboratory at UC Berkeley, supported by NIH S10 Instrumentation Grants S10RR029668 and S10RR027303. O.S.V. was supported by the Simons Foundation at the Life Sciences Research Foundation postdoctoral fellowship.

## CONTRIBUTIONS

O.S.V. and A.P.A. designed the research. O.S.V. and M.T. carried out the experiments. O.S.V. performed the computational modeling. O.S.V., M.T. and A.P.A. discussed data analyses and O.S.V. and M.T. performed the analyses. O.S.V. wrote the manuscript and O.S.V., M.T. and A.P.A. contributed to revising manuscript. S.B. performed gluconate measurements and L.J.G.C. and C.J.P. implemented shotgun proteomics.

